# An LC-MS/MS method for the quantification of tobacco-specific carcinogen protein adducts

**DOI:** 10.1101/2025.04.02.646861

**Authors:** Breanne Freeman, Chengguo Xing

**Author notes:** **Corresponding Author** Chengguo Xing – Department of Medicinal Chemistry, Center for Natural Products, Drug Discovery and Development (CNPD3), College of Pharmacy, University of Florida, Gainesville, Florida 32610, United States.

## Abstract

4-(Methyl-nitrosamino)-1-(3-pyridyl)-1-butanone (NNK) and its major metabolite 4-(methylnitrosamino)-l-(3-pyri-dine)-l-butanol (NNAL) are tobacco-specific lung carcinogens. Methods have been developed to quantify NNK- and NNAL-specific DNA adducts in pre-clinical samples but are less feasible to translation due to limited access to target tissues for sufficient DNA. NNAL-specific DNA or protein adducts have never been detected in clinical samples, which are critical to support the physiological relevance of NNAL bioactivation and carcinogenesis. We reported a sensitive and specific LC-MS/MS method to quantify hydrolyzed NNAL adduct, 1-(3-pyridinyl)-1,4-butanediol (PBD). The method was applicable to variety of biological samples, to assess tobacco exposure and estimate NNAL bioactivation.

## INTRODUCTION

Tobacco use is the leading cause for lung cancer.^1-3^ It is critical to understand tobacco carcinogenesis in humans and develop risk biomarkers to help improve lung cancer management. One well accepted root-causing mechanism of tobacco-induced tumorigenesis is through carcinogenic compounds inducing DNA damage.^5^ Tobacco specific 4-(methyl-nitrosamino)-1-(3-pyridyl)-1-butanone (NNK) and its major biometabolite 4-(methylnitrosamino)-l-(3-pyridine)-l-butanol (NNAL) have been considered as potent carcinogens with an important role in lung carcinogenesis among smokers.^7^ NNK and NNAL induce carcinogenesis through cytochrome P450 enzyme-catalyzed α-hydroxylation to produce either α-methylhydroxylation or α-methylenehydroxylation reactive intermediates that can form adducts with DNA and proteins (**Figure 1**).^8^ In addition to the common methyl adducts, NNK and NNAL respectively form adducts through pyridyloxobutylation (POB) or pyridylhydroxobutylation (PHB).^12, 13^ These DNA adducts increase the likelihood of mutations and tumorigenesis.^9^ Methods thus have been developed by Hecht and his colleagues to quantify these different types of DNA adducts.^10, 11^ Due to their low abundance and limited availability of clinical samples, these analyses have not been applicable in the clinical settings. Hecht et al later developed an alternative method by hydrolyzing the POB adducts to release 4-hydroxy-1-(3-pyridyl)-1-butanone (HPB) with successful detection of DNA adducts in buccal cells from human smokers.^15,16^ Similarly HPB has been used to detect POB protein adducts in human haemoglobin^18^ and albumin.^19^ Our recent pre-clinical work showed that HPB released from proteins positively correlated with POB DNA adducts, supporting its role as a surrogate measurement.^14^ At the same time, the hydrolyzed product from NNAL-specific PHB adducts, 1-(3-pyridinyl)-1,4-butanediol (PBD), has not been thoroughly characterized, except for our recent analyses of preclinical mouse samples with an 8-µmol NNK i.p. injection,^14^ which has minimal physiological relevance to human tobacco smoke exposure (unpublished work). To facilitate its clinical evaluation, this study developed a sensitive LC-MS/MS method to quantify PBD in a range of biological samples with different NNK exposure to assess NNAL bioactivation.

**Figure 1.**
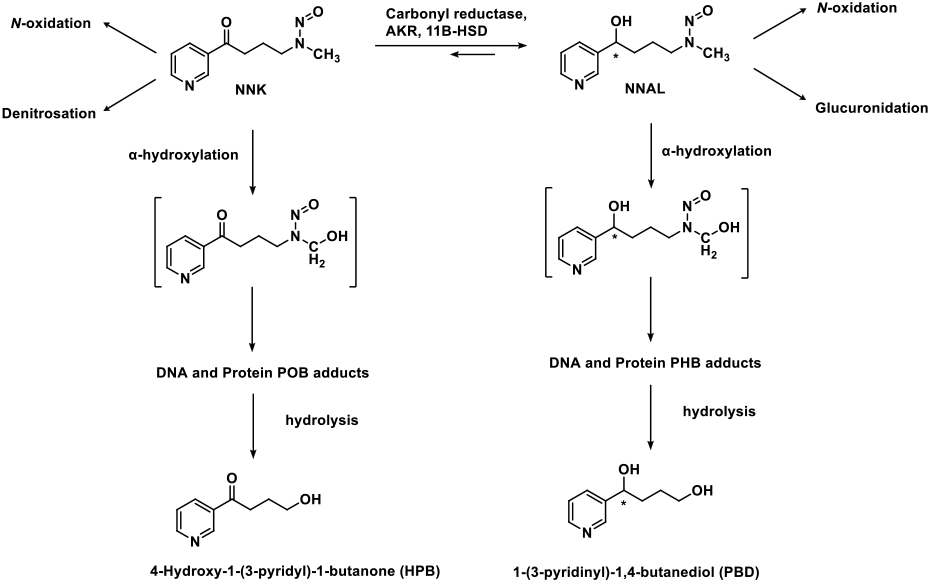
NNK and NNAL bioactivation pathway.

## EXPERIMENTAL PROCEDURES

### Caution

NNK and tobacco smoke are Class IA human carcinogens and should be handled carefully in well-ventilated fume hoods with proper protective clothing.

### Chemicals, Reagents, and Animal Diets

4-Methylnitrosamino-1-3-pyridyl-1-butanone (NNK), 4-(methylnitrosamino)-1-(3-pyridyl-[^2^H_4_])-1-butanone (NNK-d_4_), and 1-(3-pyridinyl)-1,4-butanediol (PBD) were purchased from Toronto Research Chemicals (Toronto, Ontario, Canada). AIN-93 G and AIN-M powdered diets were purchased from Harlan Teklad (Madison, WI). Liquid chromatography-mass spectrometry (LC-MS) grade of water, formic acid, methanol, ethyl acetate, and acetonitrile were purchased from Fisher Scientific (Fair Lawn, NJ). Halt Protease Inhibitor Cocktail (100x) and Pierce Universal Nuclease (250 U/µL) were purchased from Thermo Fisher Scientific (Waltham, MA).

### Tobacco Smoke Exposure Mouse Study

Female A/J mice (5-6 weeks, 16-18g) were purchased from Jackson Laboratory (Bar Harbor, ME) and maintained in the specific pathogen-free facilities, according to animal welfare protocol approved by Institutional Animal Care and Use Committee at the University of Florida (IACUC #202300000018). After one week of acclimatization, mice were randomized into two groups (n=5) and maintained on AIN-M diet. Mice were exposed to either filtered air or mainstream tobacco smoke from research cigarettes (1R6F; University of Kentucky, Lexington, NY, USA) via a SCIREQ inExpose system according to the Federal Trade Commission (FTC) protocol with a 2-h exposure daily. The cigarettes were burned at a rate of 3 cigarettes/10 minutes with target parameters of total suspended particulates (TSP) of ∼100 mg/m^3^ and CO levels of 500-600 ppm. Mice were euthanized two weeks since the first exposure and 24 hours after the last exposure via CO_2_ asphyxiation. Serum, buccal tissues, lung, and liver tissues were collected. The samples were snap frozen in liquid nitrogen and stored at -80°C until analyses.

### Mouse Protein PBD Isolation

PBD was isolated with methods slightly modified from previously established protocols.^14^ Mouse lung, liver and buccal tissues (25-35 mg) were used to isolate proteins following a previously published protocol.^14^ Isolated proteins (2 mg) were then used following the same protocol as mouse serum. Mouse serum (75 µL) protein was precipitated using 1 mL of cold methanol and the supernatant was removed. This step was repeated twice to remove free HPB and PBD as well as free NNK and NNAL. Internals standard NNK-d_4_ (2 pg in 10 µL of 0.2 pg/µL stock solution) was added to the protein samples. The protein samples were then maintained in 0.1 N NaOH (100 µL) at 37°C overnight. The samples then underwent liquid/liquid extraction using ethyl acetate (1 mL). The organic phase was collected and concentrated. The dried samples were resuspended in 30 µL of 10 mM ammonium acetate in water with 25 µL analyzed by LC-MS/MS.

### Human Protein PBD Isolation

Human PBD was isolated following the same protocol used for mouse protein isolation with using lung tissue amount of 75-100 mg and 3-4 mg of isolated protein. Plasma (150 μL) protein was precipitated using 1 mL of cold methanol and the supernatant was removed after centrifugation for 20 minutes at 13.3 kG. This step was repeated twice to remove potential free HPB and PBD as well as free NNK and NNAL. Internal standard NNK-d_4_ (2 pg in 10 μL of 0.2 pg/μL stock solution) was added to the protein samples. The samples were maintained in 0.1 N NaOH (200 μL) at 37°C overnight. The samples then underwent liquid/liquid extraction using ethyl acetate (1 mL). The organic phase was collected and concentrated. The dried samples were resuspended in 30 µL of 10 mM ammonium acetate in water with 25 µL analyzed by LC-MS/MS.

### LC-MS/MS Quantification of PBD

PBD from mice and human samples was assessed by targeted UPLC-MS^2^ consisting of a ThermoFisher Vanquish UPLC and a ThermoFisher Altis Plus triple quadrupole. The samples were resolved through an Atlantis C18 column (50 × 2.1 mm, 3 µm) with a 10.5-minute gradient with A solvent (2 mM ammonium acetate in 95% H_2_O/Acetonitrile) and B solvent (2 mM ammonium acetate in 10% H_2_O/Acetonitrile) using a 0.250 mL/min flow rate. The column temperature was held at 30°C and the autosampler was held at 5°C. PBD was resolved with a gradient starting at 1% B for 30 seconds followed by an increase to 30% B over 4 minutes. Then %B increased to 85% B over the next minute. The percentage was then held at 85% for an additional 2 minutes before returning to 1% B for the remainder of the time to equilibrate. The mass spectrometer operated in positive ionization mode. The HESI conditions were as follows, spray voltage 4000 V, sheath gas 50, aux gas 10, sweep gas 2, ion transfer tube temperature 325°C, and vaporization temperature at 350°C. The detection of the ions was performed in single reaction monitoring (SRM) mode, and the transition of PBD (168.1/117.1 m/z; CE: 15) and NNK-d_4_ (212.2/126.1 m/z; CE: 15).

## RESULTS AND DISCUSSION

### LC-MS/MS Method Development

PBD was first isolated following our previously published method used for the isolation of HPB, which uses solid phase extraction (SPE).^14^ We however observed a very low recovery of PBD through SPE (**Figure S1**). We later found a simple ethyl acetate extraction could effectively recover PBD for LCMS/MS analysis. We initially intended to use PBD-d4 as the internal standard. Although the low level of crosstalk from PBD-d4 to natural PBD was tolerable for pre-clinical samples with high levels of NNK exposure, such as i.p. NNK injection (**Figure S2**), it was problematic when analyzing samples obtained from mice with tobacco smoke exposure and human smoker samples. We therefore used NNK-d4 as the internal standard to avoid crosstalk interference.

### Calibration Curve

An eight-point curve was developed by spiking PBD into pooled nonsmoker plasma samples starting at 0.23 fmol and ending at 30 fmol (**Figure 2**). Limit of detection (LOD) and limit of quantification (LOQ) were estimated by 3.3σ/s (0.14 fmol) and 10σ/s (0.41 fmol), respectively (σ is the standard deviation of the slope (s) of the calibration curve).

**Figure 2.**
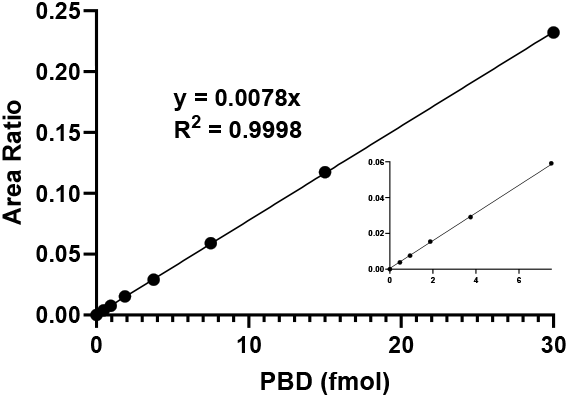
PBD Standard Curve

### Method Validation

The method was validated through assessing interday and intraday variation of three quality control samples in nonsmoker plasma. We utilized three different concentrations reflecting LOQ low concentration (0.75 fmol), middle concentration (3.0 fmol), and high concentration (24 fmol). Accuracy ranges were 95.8%-102.6% (**Table 1**). The intraday and interday variations were 1.9-5.0% and 3.4-6.7% respectively. These results demonstrate our analytical method and sample processing method are robust and reliable. We also evaluated the stability of PBD under various conditions that reflect the sample processing to determine the reliability of PBD throughout the method. We tested the three QC concentrations at autosampler (4°C), benchtop (room temperature), and hydrolysis incubation (37°C) for 24 hours. We observed an accuracy range 98.6-100.6%, 95.6-100.7%, and 101.5-104.3% respectively (**Table 2**). These results demonstrate that PBD is stable throughout sample processing and does not need special considerations.

**Table 1.**
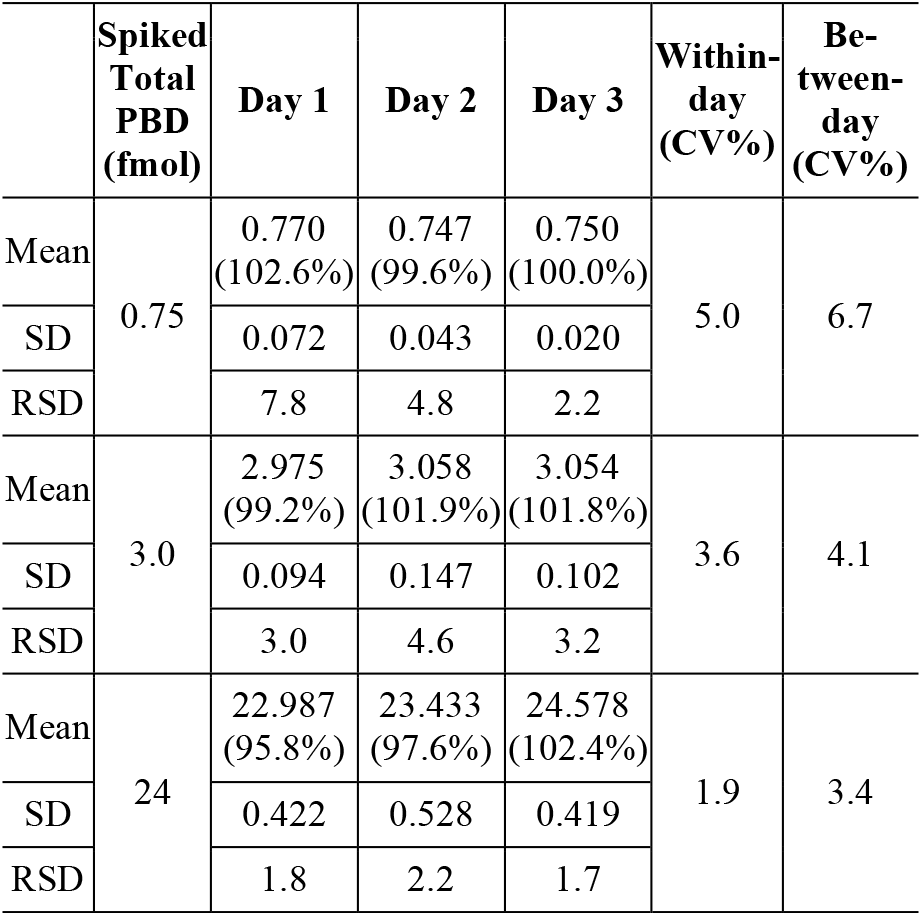
Intraday and interday precisions of PBD.

**Table 2.** Stability of PBD under different conditions.

|  | Autosampler<br>(4°C, 24h) |  |  | Benchtop<br>(rt, 24h) |  |  | Hydrolysis Incuba-<br>tion<br>(37°C, 24h) |  |  |
| --- | --- | --- | --- | --- | --- | --- | --- | --- | --- |
| Spik<br>ed<br>To-<br>tal<br>(fmo<br>l) | 0.75 | 3 | 24 | 0.75 | 3 | 24 | 0.75 | 3 | 24 |
| Mea<br>n<br>(Ac-<br>cu-<br>ra-<br>cy<br>%) | 0.754<br>(100.6<br>%) | 2.957<br>(98.6<br>%) | 23.86<br>0<br>(99.4<br>%) | 0.717<br>(95.6<br>%) | 2.892<br>(96.4<br>%) | 24.177<br>(100.7<br>%) | 0.761<br>(101.5<br>%) | 3.110<br>(103.7<br>%) | 25.034<br>(104.3<br>%) |
| SD | 0.029 | 0.028 | 0.517 | 0.010 | 0.017 | 0.446 | 0.024 | 0.026 | 0.239 |
| RSD<br>(%) | 3.2 | 0.9 | 2.2 | 1.2 | 0.5 | 1.8 | 2.6 | 0.8 | 1.0 |

### Mass Spectrometric Characterization of PBD in Mouse and Human Samples with Tobacco Smoke Exposure

We next sought to test whether PBD could be detected in physiologically more relevant tobacco smoke pre-clinical models and clinical samples. We analyzed mouse lung and buccal cell proteins with a 2-week tobacco smoke expose and were able to detect PBD in all tobacco smoke exposed samples (n=3/group), which were not detectable in the corresponding tissue samples from mice without tobacco smoke exposure (**Figure 3 A, B, C and D**). We also observe PBD in smoker plasma samples, which was absent in the plasma sample from nonsmokers (**Figure 4 A and B**). Similarly, we detected PBD in two of ten human lung tissues (**Figure 4 C and D**). These data support that our method is highly sensitive and specific to detect low levels of PBD in clinically relevant tissues to explore the potential of NNAL in human lung carcinogenesis.

**Figure 3.**
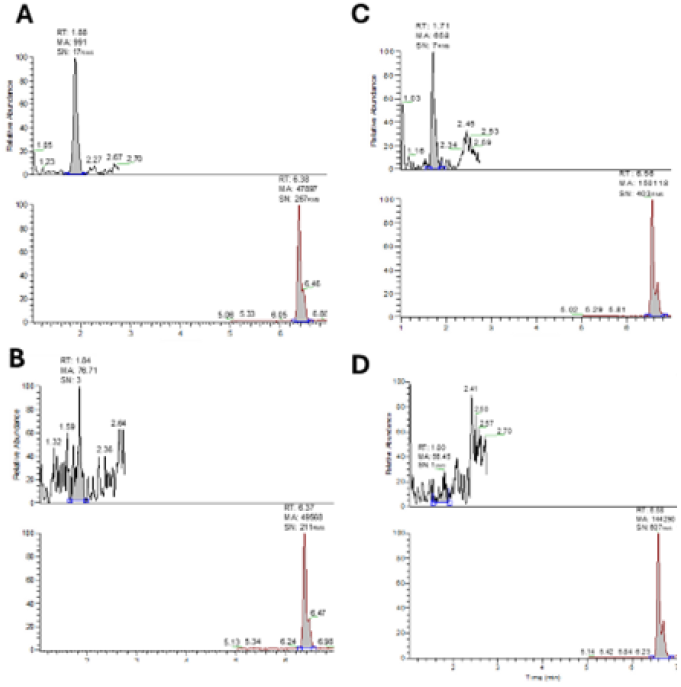
Example chromatograms representing PBD in mouse lung (A) tobacco smoke exposed, (B) not exposed, and buccal mucosa (C) exposed (D) no exposed.

**Figure 4.**
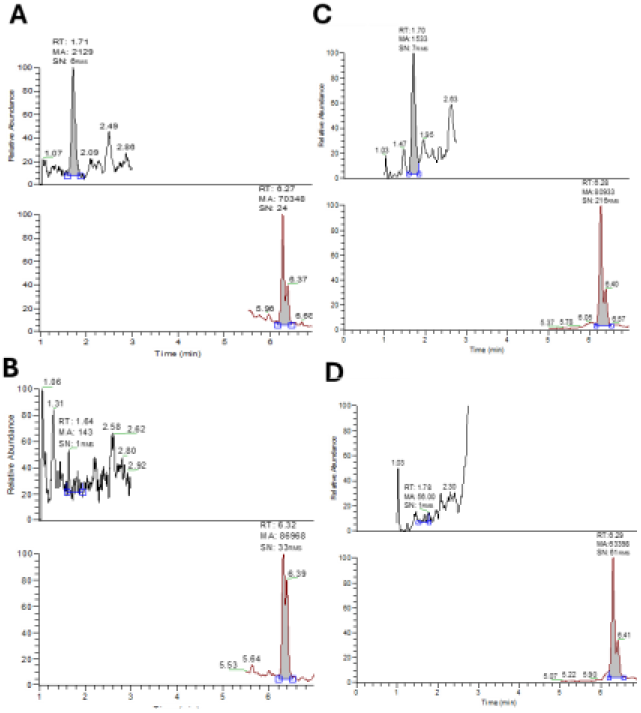
Example chromatograms representing PBD in human plasma (A) smoker (B) non-smoker and lung (C) smoker (D) non-smoker.

## CONCLUSIONS

Herein we reported a sensitive and robust LC-MS/MS assay to assess a wide range of biological samples with different to-bacco exposure that can potentially estimate the levels of NNAL bioactivation, which could be critical for tobacco smoke-induced lung cancer risk/carcinogenesis. This assay has the potential to be used to probe the relevance of NNAL bioactivation to lung carcinogenesis as it has never been studied in the context of physiologically relevant animal models or characterized in humans.

## ASSOCIATED CONTENT

### Supporting Information

The Supporting Information is available free of charge on the ACS Publications website.

Figure S1, Example chromatogram of PBD recovery with different sample preparation methods; Figure S2, Example chromatogram of internal standard crosstalk (PDF)

## AUTHOR INFORMATION

### Corresponding Author

Chengguo Xing – Department of Medicinal Chemistry, Center for Natural Products, Drug Discovery and Development (CNPD3), College of Pharmacy, University of Florida, Gainesville, Florida 32610, United States;

### Author Contributions

The manuscript was written through contributions of all authors. All authors have given approval to the final version of the manuscript. B.F. was responsible for methodology, investigation, data curation, writing, and editing of the draft. C.X. was responsible for funding acquisition, project administration, resource supervision, writing, reviewing, and editing of the draft.

## ACKNOWLEDGMENT

Research reported in this publication was partly supported by the Florida Department of Health (23B02, CX).and the National Cancer Institute of the National Institutes of Health under Award Number T32CA257923 and the UF Health Cancer Center, supported in part by state appropriations provided in Fla. Stat. § 381.915 and the National Cancer Institute of the National Institutes of Health under Award Number P30CA247796 (B.F.). The content is solely the responsibility of the authors and does not necessarily represent the official views of the National Institutes of Health or other funding sources.

---

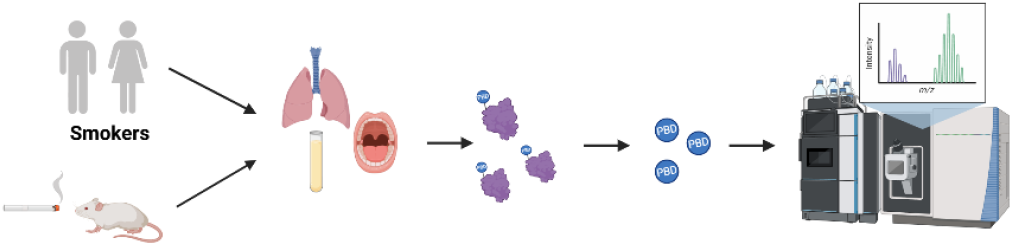

