## Supplemental Figure 1 & 2 for "An LC-MS/MS method for the quantification of tobacco-specific carcinogen protein adducts"

**Table of Contents:**

**Figure S1**. Example chromatograms of PBD recovery with different sample preparation methods.

**Figure S2**. Example chromatograms of internal standard crosstalk.

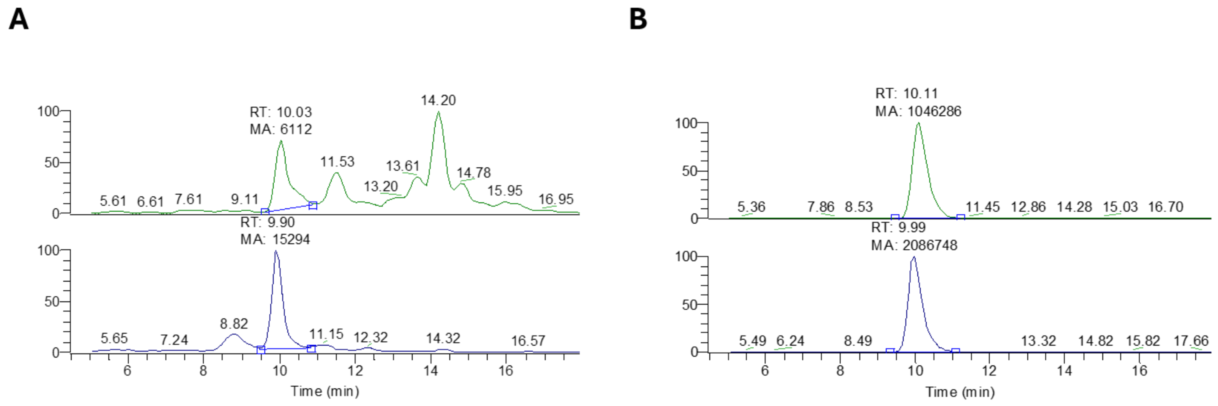


**Figure S1.** Example chromatograms of PBD signal using previously published method (**A**) and method described in this work (**B**). Quality control sample was used with PBD spiked at (25 pg total) and PBD-$d_4$ spiked at (50 pg total).

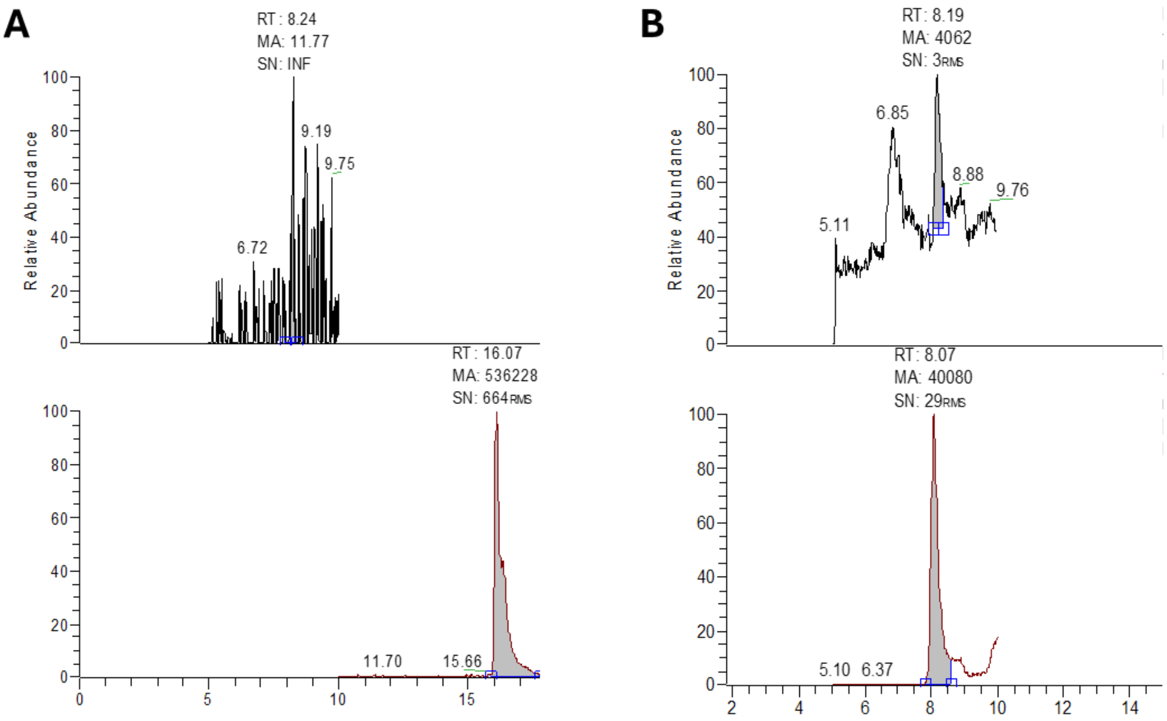


**Figure S2**. Example chromatogram of internal standard crosstalk in (**A**) NNK-d4 or (**B**) PBD-d4.
